# A new Defective Helper RNA to produce recombinant Sindbis virus that infects neurons but does not propagate

**DOI:** 10.1101/033738

**Authors:** Justus M Kebschull, Pedro Garcia da Silva, Anthony M Zador

**Author notes:** Correspondence: Anthony M Zador.

## Abstract

Recombinant Sindbis viruses are important tools in neuroscience because they combine rapid and high transgene expression with a capacity to carry large transgenes. Currently, two packaging systems based on the DH(26S)5’SIN and the DH-BB(tRNA;TE12) Defective Helper (DH) RNAs are available for making recombinant Sindbis virus that is neurotropic (able to infect neurons and potentially other cells). Both systems produce a fraction of viral particles that can propagate beyond the primary infected neuron. When injected into mouse brains, viruses produced using these DH RNAs label neurons at the injection site, but also elsewhere in the brain. Such ectopic labeling caused recombinant Sindbis viruses to be classified as anterograde viruses with limited retrograde spread, and can complicate the interpretation of neuroanatomical and other experiments.

Here we describe a new DH RNA, DH-BB(5’SIN;TE12ORF), that can be used to produce virus that is both neurotropic and propagation-incompetent. We show in mice that DH-BB(5’SIN;TE12ORF)- packaged virus eliminates infection of cells outside the injection site. We also provide evidence that ectopically labeled cells observed in previous experiments with recombinant Sindbis virus resulted from secondary infection by propagation-competent virus, rather than from inefficient retrograde spread.

Virus produced with our new packaging system retains all the advantages of previous recombinant Sindbis viruses, but minimizes the risks of confounding results with unwanted ectopic labeling. It should therefore be considered in future studies in which a neurotropic, recombinant Sindbis virus is needed.

## 1 Introduction

Sindbis virus is an enveloped, positive strand RNA virus from the family of the togaviridae. Its 11703 nucleotide genome contains two open reading frames (ORF), one coding for the nonstructural proteins, the other for the structural proteins (Fig 1a top) (Strauss et al., 1984)

**Figure 1.**
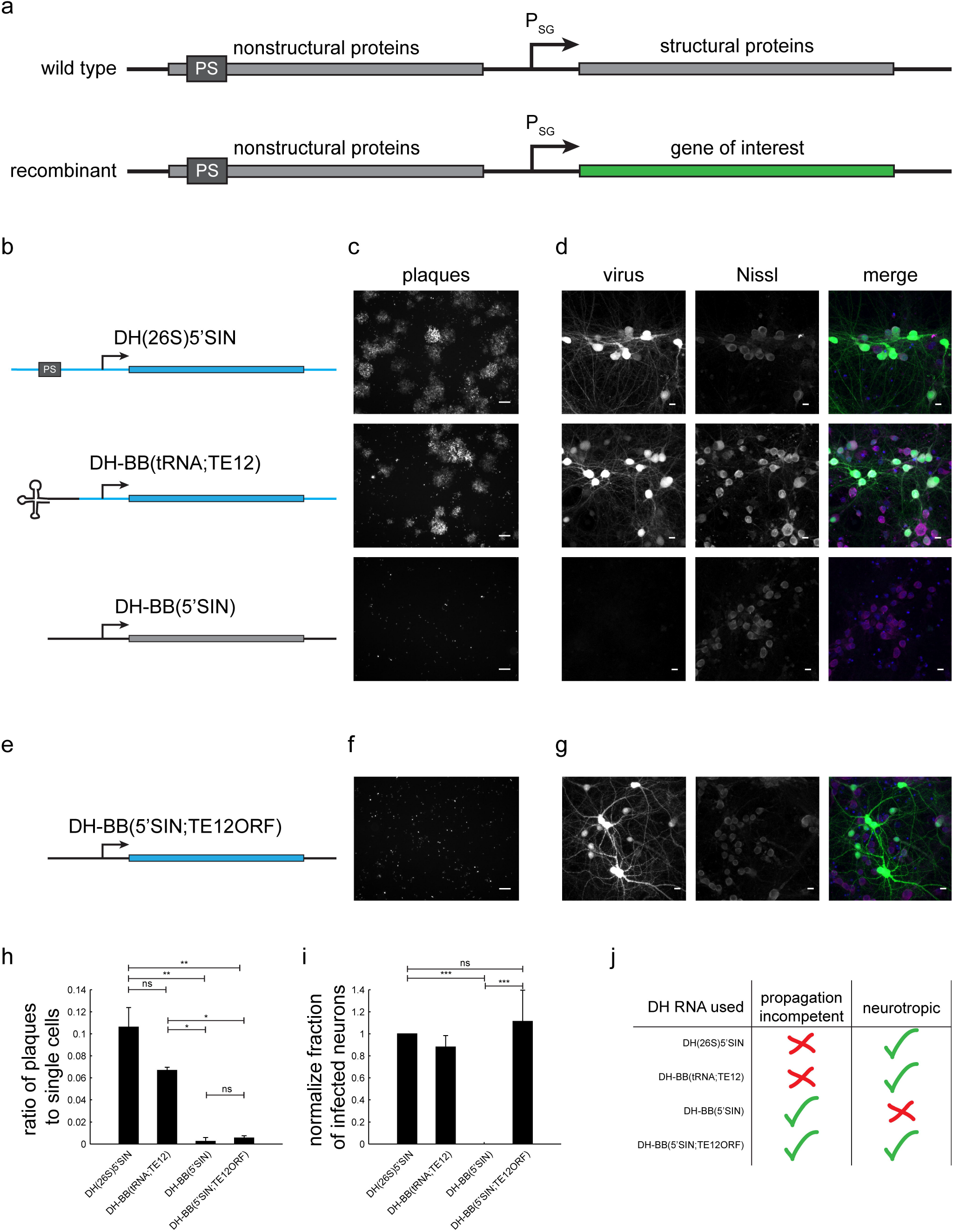
Different DH RNAs and their respective propagation competencies and neurotropism. (a) Overview of wild type and recombinant Sindbis virus genomes. (b) DH RNAs differ in their 5’ end and in the origin of their structural protein-coding region. (c) Whereas both DH(26S)5’SIN and DH-BB(tRNA;TE12) packaged virus produces GFP positive plaques in BHK cells, DH-BB(5’SIN) packaged virus does not produce plaques. Only individually infected cells are discernible. Scale bar = 100μm. (d) DH(26S)5’SIN and DH-BB(tRNA;TE12) packaged virus efficiently infect cultured hippocampal mouse neurons, while DH-BB(5’SIN) does not. Scale bar = 10μm. (e) DH-BB(5’SIN;TE12ORF) has the same structure as DH-BB(5’SIN) but carries the same structural protein open reading frame as DH(26S)5’SIN. Virus produced using DH-BB(5’SIN;TE12ORF) does not produce plaques (f) but does infect neurons (g). Quantification of the ratio of plaques to single cells (h) and the normalized fraction of infected neurons (i) shows that DH-BB(5’SIN;TE12ORF) produced virus is significantly less propagation competent than, but does infect neurons as well as DH(26S)5’SIN produced virus (mean +/- s.d.; one way ANOVA followed by Bonferroni post-hoc testing; n = 2 for plaques and n = 3 for neurons; * p-value ≤ 0.05, ** p-value ≤ 0.01, *** p-value ≤ 0.001). (j) Overview of the properties of viruses produced with the different DH RNAs.

In recombinant Sindbis vectors, the structural protein ORF is replaced with the gene of interest (Fig 1a bottom) (Xiong et al., 1989). Upon infection, the recombinant virus expresses the gene of interest instead of the structural proteins, and thus becomes a vehicle for transgene expression. Because the region encoding the essential structural proteins has been deleted, a cell infected by only recombinant Sindbis does not produce infectious viral particles. Thus to produce virus from a recombinant genome, packaging systems have been developed in which the structural proteins are supplied in *trans* by a second RNA, the DH RNA (Bredenbeek et al., 1993).

There are three classes of DH RNAs. In the first class, derived from the wild type Sindbis genome, the nonstructural protein coding region has been deleted using convenient restriction enzymes (Bredenbeek et al., 1993). The second class is similar to the first, but carries a tRNA sequence at its 5’ end rather than the usual 5’ sequence of Sindbis virus; this tRNA sequence is also found in certain naturally occurring defective interfering Sindbis particles (Bredenbeek et al., 1993). In the third class of DH RNAs, derived from naturally occurring defective interfering particles, the structural protein coding region has been reinserted (Bredenbeek et al., 1993; Geigenmüller-Gnirke et al., 1991).

A recombinant Sindbis genome does not encode all the information necessary to produce infectious viral particles. However, propagation-competent viral particles can emerge if the DH RNA is copackaged with the genome into a single particle. If this occurs, Sindbis functions effectively as a bipartite RNA virus, where the genome and DH RNA complement each other to expresses the nonstructural and structural proteins (Geigenmüller-Gnirke et al., 1991). The extent of propagation competence of recombinant Sindbis virus then depends on the rate of co-packaging of the genome with the DH RNA. Packaging of the genomic RNA is always favored over the DH RNA (Bredenbeek et al., 1993; Geigenmüller-Gnirke et al., 1991), but presence of the Sindbis packaging signal (Frolova et al., 1997) or a 5’ tRNA sequence in the DH RNA increases the rate of copackaging, and thus of the propagation competence of the resulting virus (Bredenbeek et al., 1993).

Sindbis virus has a remarkably wide host range, infecting insects and many species of higher vertebrates and many different cell types. Nonetheless, different Sindbis strains have different cell tropisms, determined by the structural proteins. Notably, the common laboratory strain Toto1101 does not infect neurons, whereas the TE12 strain does (Lustig et al., 1988). Different DH constructs are derived from different strains, and therefore produce viruses with different tropisms (Bredenbeek et al., 1993; Lustig et al., 1988).

The DH RNA conventionally used in neuroscience for Sindbis virus production is the interfering particle based DH(26S)5’SIN vector (Bredenbeek et al., 1993; Ehrengruber et al., 1999; Geigenmüller-Gnirke et al., 1991; Malinow et al., 2010). It expresses the TE12 structural proteins, making the resulting virus neurotropic (Bredenbeek et al., 1993; Lustig et al., 1988). Importantly, it also contains the Sindbis packaging signal, and produces virus that contains a fraction of propagation-competent virions (Bredenbeek et al., 1993). Previous studies have reported that, when injected into mouse brains, Sindbis virus prepared with DH(26S)5’SIN labels cells far away from the injection site, and have interpreted this as evidence for retrograde labeling (Furuta et al., 2001) (where retrograde labeling is defined as infection of the axon, and labeling of cell bodies, but no trans-synaptic spread). Despite this unintended and potentially confusing ectopic labeling, viruses produced using this DH construct have been extensively used in anterograde tracing experiments (Ghosh et al., 2011) (where anterograde labeling is defined as infection of the somatodendritic compartment and labeling of cell bodies and axons, but no trans-synaptic spread) and for transgene expression (Hayashi, 2000; Malinow et al., 2010; Shi et al., 2001).

A second neurotropic DH system was developed more recently by replacing the structural proteins in the DH-BB(tRNA) construct (Bredenbeek et al., 1993), which were derived from the non-neurotropic Toto1101 strain, with the neurotropic TE12 structural proteins from DH(26S)5’SIN (Kim et al., 2004). This change resulted in the production of neurotropic virus from the new DH-BB(tRNA;TE12) construct. However, a fraction of virions packaged with the original DH-BB(tRNA) were propagation competent (Bredenbeek et al., 1993), and the addition of TE12 structural proteins to form DH-BB(tRNA;TE12) did not appear to influence the rate of co-packaging of the DH construct. Thus, a fraction of virions produced with DH-BB(tRNA;TE12) are propagation competent.

Here we present a new Defective Helper RNA, DH-BB(5’SIN;TE12ORF), which produces a propagation-incompetent, neurotropic virus. When injected into mouse brains, virus produced with this new DH RNA only very rarely labels neurons outside the injection site, suggesting that the ectopic labels observed for Sindbis viruses derived from other DH RNAs arose from secondary infection by propagation-competent particles. Our new DH RNA therefore provides precise spatial control of Sindbis virus-based expression in the mouse brain, and removes ectopic infection as a confounding factor in neuroanatomical and physiological experiments.

## 2 Results and Discussion

### 2.1 Generation of a propagation-incompetent neurotropic Sindbis packaging system

To investigate the replication competence and neurotropism of different Sindbis packaging systems, we generated a recombinant genome that expressed GFP, and packaged it with three different DH RNAs (Fig 1b): (i) the commonly used neurotropic DH(26S)5’SIN (Bredenbeek et al., 1993), (ii) the neurotropic DH-BB(tRNA;TE12) (Kim et al., 2004) and (iii) the Toto1101 derived DH-BB(5’SIN) (Bredenbeek et al., 1993) (Fig 1b). As expected (Bredenbeek et al., 1993; Kim et al., 2004), plaque assays in BHK cells revealed that both DH(26S)5’SIN and DH-BB(tRNA;TE12) produced propagation-competent virus. In contrast, DH-BB(5’SIN)-derived virus showed no evidence of replication competence (Fig 1c), in accordance with the lack of packaging signals and 5’ terminal tRNA sequences in the DH RNA. Conversely, the propagation-competent DH(26S)5’SIN and DH-BB(tRNA;TE12)-derived viruses infected primary mouse hippocampal cultures, whereas propagation-incompetent DH-BB(5’SIN)-derived virus failed to infect neurons (Fig1d) (Bredenbeek et al., 1993; Lustig et al., 1988).

We sought to combine the propagation-incompetence of the DH-BB(5’SIN) system with the neurotropism provided by the other two DH RNAs. Tropism is controlled by the structural proteins (Lustig et al., 1988), replication competence by the probability of co-packaging of the helper RNA with the genomic RNA (Bredenbeek et al., 1993; Geigenmüller-Gnirke et al., 1991). We therefore replaced only the structural protein ORF of the DH-BB(5’SIN) vector with the structural protein ORF of the DH-BB(tRNA;TE12) construct (Fig 1e). As predicted, virus produced from the resulting DH-BB(5’SIN;TE12ORF) construct does not show evidence of secondary infection in a plaque assay (Fig 1f), but does infect neurons (Fig 1g). Quantification of the propagation incompetence (Fig 1h) and neurotropism (Fig 1i) of viruses produced using the different DH RNAs confirms that DH-BB(5’SIN;TE12ORF) maintained the propagation incompetence of DH-BB(5’SIN) and the neurotropism of DH-BB(tRNA;TE12) producing a combination of propagation incompetence and neurotropism previously not available in recombinant Sindbis viruses (Fig 1j).

### 2.2 Use of DH-BB(5’SIN;TE12ORF) eliminates labeling of cells outside the injection site

We next characterized the behavior of viruses packaged with DH(26S)5’SIN and DH-BB(5’SIN;TE12ORF) *in vivo.* To do so, we injected equal amounts of virus (as determined by qPCR titering; see Materials and Methods) into the locus coeruleus of 3 mice per virus and examined the pattern of infection.

Virus packaged with DH(26S)5’SIN efficiently infected neurons at the primary injection site (Fig 2a inset). It also labeled numerous cells spread out across the brain far away from the injection site (228 +/- 166 cells per animal (mean +/- s.d.); Fig 2a). Such spread has previously been reported for virus packaged with DH(26S)5’SIN, and was attributed to some low level of retrograde infection (Furuta et al., 2001). However, close inspection of those putatively retrogradely labeled cells revealed GFP expression not only in isolated projection neurons with long axons that might have reached the site of injection (Fig 2b), but also in interneurons whose axons remain local and do not extend to the injection site and therefore cannot be labeled by retrograde infection (Fig 2c). Furthermore, we also observed clusters of up to 50 GFP-expressing neurons (Fig 2d). Such dense labeling is again difficult to explain by inefficient retrograde infection, which inherently favors sparse labeling over large areas, but would be expected from occasional propagation of infective particles from the axons of neurons infected at the injection site.

**Figure 2.**
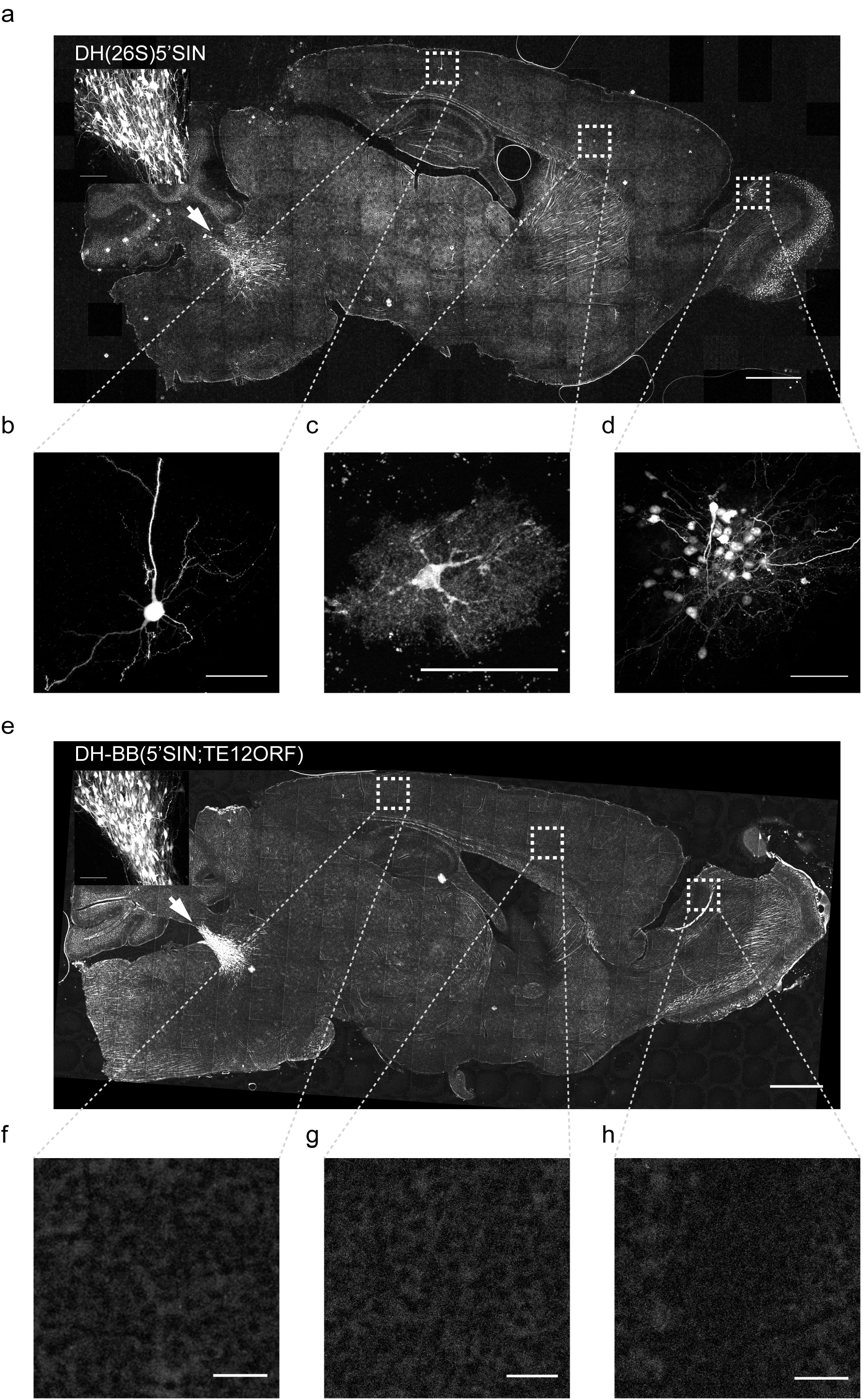
Virus produced with DH-BB(5’SIN;TE12ORF) does not label cells outside the primary injection site. (a) Overview of a representative section of a mouse brain injected with DH(26S)5’SIN packaged virus into locus coeruleus. Arrow labels the injection site; scale bar = 1mm. The inset shows the center of the injection site. Scale bar = 100μm. Ectopically labeled cells, such as a pyramidal neuron in cortex (b), but also an interneuron in cortex (c) and a cluster of local granule cells in the olfactory bulb (d) are discernible. Scale bar = 50μm. (e) Overview of a representative section of a mouse brain injected with DH-B(5’SIN;TE12ORF) packaged virus into locus coeruleus. Arrow labels the injection site; scale bar = 1mm. The inset shows the center of the injection site. Scale bar = 100μm. (f-h) No ectopically labeled cells can be detected. Scale bar = 50μm. Bright spots in the overview image were ruled out as ectopically labeled cells as they are often much larger than cells and are lacking any cellular morphology. They instead appear to be the result of brightly autofluorescent contaminants in the mounting media.

Virus packaged with our newly engineered DH-BB(5’SIN;TE12ORF) packaging system labeled neurons at the injection site with an efficiency similar to that of DH(26S)5’SIN (Fig 2e inset). However, in contrast to DH(26S)5’SIN-packaged virus, virus packaged with DH-BB(5’SIN;TE12ORF) showed almost no ectopic infection (Fig 2e-h). In a total of 3 mice, we detected only a single labeled cell outside the injection site, compared with hundreds of cells per animal using the previous system (p = 0.039; Student’s t-test). Thus use of DH-BB(5’SIN;TE12ORF) eliminates the co-packaging of DH RNA, and thereby eliminates the secondary infection responsible for ectopic spread beyond the injection site. Accordingly, we classify recombinant Sindbis virus as purely anterograde viruses in the mouse brain.

## 3 Conclusion

We here describe a new Sindbis virus packaging system that produces a neurotropic virus that does not propagate beyond *in vitro* or *in vivo.* Secondary infection is hard to control and can have unintended effects in neuroanatomical and physiological studies. Our new packaging system should therefore be used in all future studies in which secondary infection can confound the interpretation of results.

## 4 Materials and Methods

### 4.1 Defective Helper constructs

The three DH constructs used in this study were generous gifts from the following laboratories: Robert Malinow (UCSD; DH(26S)5’SIN), Jinny Kim (KIST; DH-BB(tRNA;TE12)) and Charles Rice (Rockefeller University; DH-BB(5’SIN)). To produce DH-BB(5’SIN;TE12ORF) we performed PCR for the TE12 ORF on DH-BB(tRNA;TE12) using forward primer 5’-TAA AGC ATC TCT ACG GTG GTC C-3’ and reverse primer 5’-CAT CGA GTT TTG CTG GTC GGA-3’ to produce a 3.8 kb fragment. We amplified the DH-BB(5’SIN) backbone excluding the structural ORF by PCR using forward primer 5’-CCG CTA CGC CCC AAT GAT CC-3’ and reverse primer 5’-TAT TAG TCA GAT GAA ATG TAC TAT GCT GAC-3’ to produce a 3 kb fragment. We performed all PCRs using Q5 High Fidelity 2x Master Mix (New England Biolabs) according to the manufacturer’s instructions using 72°C annealing temperatures. We then assembled the two PCR fragments using Gibson assembly master mix (New England Biolabs) according to the manufacturer’s protocol and transformed chemically competent TOP10 cells with the assembly mix. The DH-BB(5’SIN;TE12ORF) plasmid is available from Addgene.org under accession 72309.

### 4.2 Sindbis virus production

To produce high-titer Sindbis virus, we first linearized both a genomic construct encoding GFP and the DH plasmid construct with PacI or XhoI (New England Biolabs), respectively. After Phenol:Chloroform:Isoamyl alcohol (Thermo Fisher) purification of the linearize plasmids, we *in vitro* transcribed genomic and DH RNAs using the mMessage mMachine SP6 *in vitro* transcription kit (Thermo Fisher) according to the manufacturer’s instructions. We then cleaned the in vitro transcribed RNAs by Acid Phenol:Chloroform (Thermo Fisher) and transfected BHK-21 cells with a 1.7:1 weight ratio of genomic to DH RNA using Lipofectamine 2000 (Thermo Fisher) in complete media (alpha-MEM, 5% heat inactivated FBS, 1% PennStrep, 1% MEM vitamins, 1% 200mM L-Glutamine (all Thermo Fisher)) according to the manufacturer’s instructions. Forty hours after transfection, we removed the supernatant and concentrated the virus by ultracentrifugation, as previously described (Malinow et al., 2010).

### 4.3 Sindbis virus titering

To obtain a measure of Sindbis titer independent of the virus’s tropism, we quantified the number of RNaseI-protected genomic RNAs per ml of virus (GC/ml). Briefly, we digested 1¼l of virus with 1U of RNaseI (Epicentre) in 50μl total volume at 37°C for 30 minutes and then extracted protected RNA using Trizol reagent (Thermo Fisher) according to the manufacturer’s protocol. We reverse transcribed the genomic RNA using the gene specific primer 5’-GGG TCG CCT TGC TTG AAG TG-3’ that falls into the nsp1-4 region of the genomic RNA using SuperScriptIII reverse transcriptase (Thermo Fisher) according to the manufacturer’s protocol. We then measured the number of cDNA molecules relative to a plasmid standard by qPCR using SYBR Green power Mastermix (Thermo Fisher) and primers 5’-TAT CCG CAG TGC GGT TCC AT-3’ and 5’-TGT CGC TGA GTC CAG TGT TGG-3’.

### 4.4 Primary hippocampal neuron culture

Animal procedures were approved by the Cold Spring Harbor Laboratory Animal Care and Use Committee and carried out in accordance with National Institutes of Health standards.

We prepared dissociated hippocampal neurons from E18 CD1 mouse pups as previously described (Pak et al., 2001) and plated them in XONA Microfluidic chambers (SND450, Xona Microfluidics) according to the manufacturer’s instructions. At 14-21 days in vitro we infected cultures with 1μl of 10^10^ genome copies/ml virus as previously described (Taylor et al., 2010). We fixed the cells in 4% paraformaldehyde (Electron Microscopy Sciences) in 0.1M phosphate buffer 24 hours after infection and stained them using Neurotrace 640/660 Deep Red Fluorescent Nissl Stain (Thermo Fisher) according to the manufacturer’s protocol. We then imaged the cells on a Zeiss LSM780 confocal microscope using a 40x oil-immersion objective and quantified the number of both Nissl positive and GFP positive cells by hand.

### 4.5 Plaque assays

We infected 90% confluent BHK-21 cells with Sindbis virus in 200μl volume for 1 hour in 24-well plates. We then overlaid the cells with 0.4% SeaPlaque Agarose (Lonza) in complete culture media. 24 hours post infection we fixed the cells in 4% paraformaldehyde in 0.1M phosphate buffer and imaged three independent regions of GFP positive cells on a Zeiss Observer microscope per biological replicate. We then quantified the ratio of GFP plaques to single infected cells using a custom ImageJ macro that distinguishes plaques from single cells based on size. We used a threshold of 1000 pixels to distinguish single cells from plaques (see Supplementary Materials for source code).

### 4.6 Virus injections

Animal procedures were approved by the Cold Spring Harbor Laboratory Animal Care and Use Committee and carried out in accordance with National Institutes of Health standards.

We pressure injected 180nl of 2 × 10^10^ GC/ml virus into the right locus coeruleus (5.4mm posterior, 0.8mm lateral, 2.9mm and 3.1mm deep from the surface of the brain; 90nl per depth) of 10-14 week old C57/BL6 males as described (Cetin et al., 2007). 48 hours post infection we transcardially perfused animals with ice cold saline (9g/l) followed by ice cold 4% paraformaldehyde (Electron Microscopy Sciences) in 0.1M phosphate buffer. We post fixed brains in 4% paraformaldehyde in 0.1M phosphate buffer overnight at 4°C and then cut them into 70μm thick sections on a Vibratome and mounted the sections in Vectashield (Vector Labs). We acquired all overview images and the detailed images of DH-BB(5’SIN;TE12) injected brains using a Olympus IX71 inverted epifluorescence microscope with a 10x dry objective. We here presented background subtracted maximum projections. We imaged the ectopically infected cells of DH(26S)5’SIN injected brains and the injection sites of both brains using a Zeiss LSM780 confocal microscope with a 40x oil-immersion and a 20x dry objective, respectively, and presented maximum projections.

### 6 Author Contributions

JK and AZ conceived the study. JK and PS performed the experiments. JK and AZ wrote the paper.

### 7 Funding

This work was supported by the following funding sources: National Institutes of Health [5RO1NS073129 to A.M.Z., 5RO1DA036913 to A.M.Z.]; Brain Research Foundation [BRF-SIA-2014-03 to A.M.Z.]; Simons Foundation [382793/SIMONS to A.M.Z.]; PhD fellowship from the Boehringer Ingelheim Fonds to J.M.K.; PhD fellowship from the Genentech Foundation to J.M.K.; PhD fellowship from the Fundaçâo para a Ciência e Tecnologia, Portugal to P.G.S.

## 8 Acknowledgments

The authors would like to acknowledge Zhifan Yang for help with cell counting and Nour El-Amine for imaging using the Deltavision microscope. This work was performed with assistance from the CSHL Microscopy Shared Resource which is supported by Cancer Center Support Grant 5P30CA045508.

## References

Bredenbeek, P. J., Frolov, I., Rice, C. M., and Schlesinger, S. (1993). Sindbis virus expression vectors: packaging of RNA replicons by using defective helper RNAs. Journal of Virology 67, 6439–6446.

Cetin, A., Komai, S., Eliava, M., Seeburg, P. H., and Osten, P. (2007). Stereotaxic gene delivery in the rodent brain. Nat Protoc 1, 3166–3173. doi:10.1038/nprot.2006.450.

Ehrengruber, M. U., Lundstrom, K., Schweitzer, C., Heuss, C., Schlesinger, S., and Gähwiler, B. H. (1999). Recombinant Semliki Forest virus and Sindbis virus efficiently infect neurons in hippocampal slice cultures. Proceedings of the National Academy of Sciences 96, 7041–7046.

Frolova, E., Frolov, I., and Schlesinger, S. (1997). Packaging signals in alphaviruses. Journal of Virology 71, 248–258.

Furuta, T., Tomioka, R., Taki, K., Nakamura, K., Tamamaki, N., and Kaneko, T. (2001). In vivo transduction of central neurons using recombinant Sindbis virus: Golgi-like labeling of dendrites and axons with membrane-targeted fluorescent proteins. J. Histochem. Cytochem. 49, 1497–1508.

Geigenmüller-Gnirke, U., Weiss, B., Wright, R., and Schlesinger, S. (1991). Complementation between Sindbis viral RNAs produces infectious particles with a bipartite genome. Proceedings of the National Academy of Sciences 88, 3253–3257.

Ghosh, S., Larson, S. D., Hefzi, H., Marnoy, Z., Cutforth, T., Dokka, K., et al. (2011). Sensory maps in the olfactory cortex defined by long-range viral tracing of single neurons. Nature 472, 217–220. doi:10.1038/nature09945.

Hayashi, Y. (2000). Driving AMPA Receptors into Synapses by LTP and CaMKII: Requirement for GluR1 and PDZ Domain Interaction. Science 287, 2262–2267. doi:10.1126/science.287.5461.2262.

Kim, J., Dittgen, T., Nimmerjahn, A., Waters, J., Pawlak, V., Helmchen, F., et al. (2004). Sindbis vector SINrep(nsP2S726): a tool for rapid heterologous expression with attenuated cytotoxicity in neurons. Journal of Neuroscience Methods 133, 81–90.

Lustig, S., Jackson, A. C., Hahn, C. S., Griffin, D. E., Strauss, E. G., and Strauss, J. H. (1988). Molecular basis of Sindbis virus neurovirulence in mice. Journal of Virology 62, 2329–2336.

Malinow, R., Hayashi, Y., Maletic-Savatic, M., Zaman, S. H., Poncer, J.-C., Shi, S.-H., et al. (2010). Introduction of green fluorescent protein (GFP) into hippocampal neurons through viral infection. Cold Spring Harbor Protocols 2010, pdb.prot5406–pdb.prot5406. doi:10.1101/pdb.prot5406.

Pak, D. T. S., Yang, S., Rudolph-Correia, S., Kim, E., and Sheng, M. (2001). Regulation of Dendritic Spine Morphology by SPAR, a PSD-95-Associated RapGAP. Neuron 31, 289–303. doi:10.1016/S0896-6273(01)00355-5.

Shi, S., Hayashi, Y., Esteban, J. A., and Malinow, R. (2001). Subunit-specific rules governing AMPA receptor trafficking to synapses in hippocampal pyramidal neurons. Cell 105, 331–343.

Strauss, E. G., Rice, C. M., and Strauss, J. H.(1984). Complete nucleotide sequence of the genomic RNA of Sindbis virus. Virology 133, 92–110.

Taylor, A. M., Dieterich, D. C., Ito, H. T., Kim, S. A., and Schuman, E. M. (2010). Microfluidic Local Perfusion Chambers for the Visualization and Manipulation of Synapses. Neuron 66, ST-68. doi:10.1016/j.neuron.2010.03.022.

Xiong, C., Levis, R., Shen, P., Schlesinger, S., Rice, C. M., and Huang, H. V. (1989). Sindbis virus: an efficient, broad host range vector for gene expression in animal cells. Science 243, 1188–1191.

